# Pregnancy-associated oxidative stress and inflammation are not associated with impaired maternal neuronal activity or memory function

**DOI:** 10.1101/2024.01.26.577461

**Authors:** Jessica L. Bradshaw, E. Nicole Wilson, Jennifer J. Gardner, Steve Mabry, Selina M. Tucker, Nataliya Rybalchenko, Edward Vera, Styliani Goulopoulou, Rebecca L. Cunningham

## Abstract

Pregnancy is associated with neural and behavioral plasticity, systemic inflammation, and oxidative stress. Yet, the impact of systemic inflammation and oxidative stress on maternal neural and behavioral plasticity during pregnancy are unclear. We hypothesized that the maternal hippocampal CA1, a brain region associated with cognition, would be protected from pregnancy-associated systemic elevations in inflammation and oxidative stress, mediating stable peripartum cognitive performance. Cognitive performance was tested using novel object recognition (recollective memory), Morris water maze (spatial memory), and open field (anxiety-like) behavior tasks in female Sprague-Dawley rats of varying reproductive states [non-pregnant (nulliparous), pregnant (near term), and two months post-pregnancy (primiparous); n = 7-8/group]. Plasma and CA1 proinflammatory cytokines were measured using a MILLIPLEX® magnetic bead assay. Plasma oxidative stress was measured via advanced oxidation protein products (AOPP) assay. CA1 markers of oxidative stress, neuronal activity, and apoptosis were quantified via western blotting. Our results demonstrate CA1 oxidative stress-associated markers were elevated in pregnant compared to nulliparous rats (*p* ≤ 0.017) but were equivalent levels in pregnant and primiparous rats. In contrast, reproductive state did not impact CA1 inflammatory cytokines, neuronal activity, or apoptosis. Likewise, there was no effect of reproductive state on recollective or spatial memory. Even so, spatial learning was impaired (*p* ≤ 0.007) while anxiety-like behavior (*p* ≤ 0.034) was reduced in primiparous rats. Overall, our data suggest maternal hippocampal CA1 is protected from systemic inflammation but vulnerable to peripartum oxidative stress. Thus, peripartum oxidative stress elevations, such as in pregnancy complications, may contribute to peripartum neural and behavioral plasticity.

## INTRODUCTION

Motherhood requires widespread structural and functional adaptations in numerous organ systems during pregnancy and post-pregnancy to ensure offspring development and suitable maternal caregiving behavior (1, 2). Specifically, the maternal central nervous system undergoes extensive remodeling during pregnancy and post-pregnancy (2, 3). Clinical evidence suggests that the total brain volume of mothers decreases during pregnancy and recovers within 6 months postpartum (3). Even so, maternal brain structural and functional dynamics have been shown to be region-specific during pregnancy and post-pregnancy (4–8), and these region-specific changes can be sustained for years following parturition (4, 5). Notably, a recent study revealed pregnancy-associated reductions in grey matter volume within regions associated with cognition and mood are sustained through two years post-pregnancy in human cohorts (5).

Even though these pregnancy-associated physiological adaptations prepare mothers for the transition to parenthood, many of these adaptations are also associated with peripartum mental health disorders and crises, including peripartum depression, anxiety, and cognitive impairments (9–14). Indeed, approximately 80% of pregnant women self-report lived experiences of cognitive impairment during pregnancy or post-pregnancy (15), and pregnancy-associated cognitive declines including impairments in learning and working memory have been identified in women during pregnancy and post-pregnancy (15–18). Moreover, women experiencing pregnancy complications such as preeclampsia, an obstetric disorder characterized by elevated systemic oxidative stress and inflammation, are at increased risk of developing cognitive impairments during pregnancy and later in life (19–21). Although there is growing evidence of structural brain fluctuations and cognitive impairments during pregnancy and post-pregnancy, the underlying mechanisms contributing to maternal neuroplasticity and behavior remain elusive.

Reproductive status can impact numerous biological factors that modulate neuroplasticity and behavior (16). For instance, pregnancy-associated temporal patterns in hormone secretion modify the maternal brain and shape maternal behavior during pregnancy, parturition, and post-pregnancy (22–24). Additionally, systemic fluctuations in the maternal immune response occur during the peripartum period and have been associated with alterations in neuroplasticity and maternal behavior (2, 25–28). Yet, few studies have examined the localized neuroimmune response and its association with systemic inflammation during pregnancy and post-pregnancy. Moreover, systemic inflammation is a prominent cause of oxidative stress, and oxidative stress impacts behavioral responses and neuronal processes including neurogenesis, migration, and synaptic pruning (29–32). Still, little is known regarding the impact of reproductive experience on inflammation and oxidative stress within brain regions associated with maternal cognitive functions.

The hippocampus and amygdala are two of the most vulnerable regions to oxidative stress and inflammatory insults, as these regions are reported as the first regions to undergo functional decline following exposure to stressors (32). Particularly, the hippocampus is a highly plastic brain region that regulates learning and memory functions, as well as the production of new neurons (33, 34). Within the hippocampus, CA1 neurons are critically involved in episodic recollective memory (35). Importantly, the hippocampus and amygdala are interconnected brain regions that cooperatively contribute to anxiety-like behavior (36). Of note, previous studies have demonstrated reductions in hippocampal volume and neurogenesis in women during pregnancy and post-pregnancy (2, 16), as well as transcriptome alterations in the amygdala following parturition (37).

In this study, we used female rats with varying reproductive experiences (nulliparous, pregnant, and primiparous) to determine the impact of reproductive experience on maternal cognitive function and investigate oxidative stress and inflammation as modulators of maternal cognitive function. We hypothesized that the maternal brain is protected from pregnancy-associated elevations in systemic inflammation and oxidative stress, mediating stable cognitive performance during pregnancy and post-pregnancy.

## MATERIALS & METHODS

### Animals

All protocols were approved by the Institutional Animal Care and Use Committee (IACUC) of the University of North Texas Health Science Center (Protocol # IACUC-2020-032). Protocols were performed in accordance with the National Institutes of Health’s *Guide for the Care and Use of Laboratory Animals*. All experiments were conducted using nulliparous (non-pregnant virgin, 9-11 weeks old on arrival) or timed-pregnant (13-15 weeks old on arrival) Sprague-Dawley female rats purchased from Envigo (Indianapolis, IN and Houston, TX). Pregnant rats arrived at the University of North Texas Health Science Center animal facilities on gestational day (GD) 10 (GD 1 = first day of sperm plug; term = 22-23 days). Male rats were excluded from these studies because the focus of the research is the effects of pregnancy history (a female-specific condition) on maternal physiology.

Female rats were single-housed upon arrival under 12-h:12-h reverse light/dark cycles (lights off 07:00 h, lights on 19:00 h) in a temperature and humidity-controlled environment. Animals were provided standard laboratory chow and water *ad libitum*. Following one week of acclimatization to the animal facilities, female rats were familiarized with operator handling to reduce stress responses during behavioral testing. This study comprised a cross-sectional design that consisted of three groups characterized by reproductive experience at time of behavioral testing and sample collection: nulliparous (NULLI, virgins), late gestation pregnancy (PREG, GD 20-21), and primiparous (PRIMI, two months following first pregnancy). Timed-pregnant rats were assigned to PREG and PRIMI groups. Rats assigned to PRIMI group were allowed to give birth, and pups were culled to six pups per litter within 48 hours of delivery. Pups were weaned at postnatal day 28. Behavior testing of the PRIMI group began one week after weaning to assess behaviors without the confounding effects of pregnancy and lactation-associated hormonal fluctuations. The total number of animals used in this study were 31 female rats assigned as NULLI (n= 16), PREG (n = 8), and PRIMI (n= 7). All experiments were performed when female rats were 3-6 months old.

### Behavioral Testing

Behavioral studies were conducted under red lighting over two days during the rodent’s active period (08:00 h to 14:00 h). PREG and PRIMI rats were tested in separate cohorts alongside NULLI rats. Behavioral tests were used to assess hippocampal-associated memory (novel object recognition and Morris water maze) and anxiety-like behavior (open field exploration). Rats were placed in carriers and acclimated to the behavior testing room for 30 minutes prior to beginning testing. All testing equipment was thoroughly cleaned with 70% ethanol between each animal. Behaviors were recorded with ANY-Maze video tracking software (Stoelting Co., version 5.14) for subsequent analysis by an investigator blinded to assigned groups.

#### Novel object recognition

Novel object recognition was performed to assess short-term memory as previously described (38). Briefly, two objects of the same color, shape, and size were placed in adjacent corners at the top of the arena [24” (W) x 24” (L) x 12” (H)]. Animals were individually placed facing away from the objects in the bottom of the arena and allowed to explore the arena and objects for five minutes. After habituation to the arena and objects, animals were returned to their respective carriers to rest for one hour. To assess short-term memory, one of the objects from the previous phase was placed back into the arena at its previous location (familiar object) while the other object was replaced in the same location with a novel object of different shape, color, and texture (novel object). After resting for one hour, the animals were allowed to explore the arena and two objects (one familiar and one novel) for three minutes. Latency to initial movement, latency to novel object and familiar object, total contacts with the novel object and familiar object, object preference, and distance traveled during the test were examined.

#### Morris water maze

To assess learning and spatial memory, the Morris water maze was performed as previously described (38, 39). Briefly, rats were trained to swim to a visible platform approximately 1 cm above the water surface of an opaque pool (23-25°C) on day 1 of the behavioral task. On days 2-4 of training (learning phase), rats were trained to locate a submerged target platform within 90 seconds (3 trials/day), and each rat was allowed 20 seconds to sit on the located platform to observe visual cues placed on the walls to aid in the formation of spatial memory. On day 5 (probe trial), the submerged target was removed and each rat was provided 30 seconds to swim to the target location and search for the platform. Learning performance on days 2-4 was assessed by quantifying the latency and pathlength to the target, as well as a learning index (40). Spatial memory was assessed during the probe trial on day 5 by quantifying latency and pathlength to the target zone.

#### Open field

Anxiety-like behaviors were assessed in an open field arena. Animals were individually placed in the bottom of the open arena [24” (W) x 24” (L) x 12” (H)] facing away from the open arena and allowed to explore for five minutes. The number of entries into the center of the open field, duration in the center, distance traveled in the center, and total distance traveled were assessed.

### Euthanasia and Tissue Harvest

Animals were anesthetized with 2-3% isoflurane and euthanized via decapitation during the animals’ active phase (09:00-11:00). All animals were euthanized within four days following the final day of behavior testing. Trunk whole blood was collected and plasma was isolated using EDTA-coated collection tubes (BD, Cat. No. 367856) centrifuged at 2,000 x g for 10 minutes at 4°C. Freshly isolated plasma was aliquoted and stored at −80°C until further analysis. Following euthanasia, brains were quickly removed, flash-frozen in 2-methylbutane (Millipore Sigma, Cat. No. MX0760), and stored at −80°C until further analysis.

### Sample Preparation

Frozen brains were thawed in chilled 1X phosphate-buffered saline (Fisher, Cat. No. BP399) and sliced into 1-mm coronal sections using a brain matrix (ASI Instruments, Cat. No. RBM-4000C). Brain nuclei within the CA1 of the dorsal hippocampus (−5.30 mm from Bregma) and basolateral and basomedial amygdala (−3.30 mm from Bregma) were microdissected according to Paxinos and Watson’s brain atlas (41) using blunt 20-gauge needles attached to 1 ml syringes. Microdissected brain punches were stored at −80°C until further analysis. Frozen CA1 and amygdala microdissections were thawed in RIPA lysis buffer (VWR, cat #N653) containing (per 0.5 ml): 2.5 µl Halt™ protease and phosphatase inhibitor (Thermo Scientific, Ca. No. 78442), 1 µl 0.5 M ethylenediaminetetraacetic acid (EDTA, Thermo Scientific, Ca. No. J15694.AE), and 1 µl 0.5 mM dithiothreitol (DTT, Millipore Sigma, Cat. No. 43815) and homogenized as previously described (42). Total protein concentrations in CA1 and amygdala homogenates were quantified using Pierce BCA Protein Assay (Thermo Scientific Cat. No. 23225).

### Plasma and Brain Cytokine Analysis

Plasma and dorsal hippocampal CA1 cytokines were assessed using a MILLIPLEX® Rat Cytokine/Chemokine Magnetic Bead Panel (Millipore Sigma Cat. No. RECYTMAG-65K) customized for detection of pro-inflammatory (TNF-α, IL-6, IL-1β, IL-17A) and anti-inflammatory (IL-10) cytokines. Plasma samples were diluted 1:2 while CA1 lysates were diluted 1:10 in assay buffer prior to running the assay according to the manufacturer’s instructions. All samples, standards, and quality controls were plated in duplicate, and cytokines were measured on a Luminex® 200 instrument using xPONENT® software version 4.3 (Luminex Corporation, Austin, TX). Quality control values for each cytokine were within the range provided by the manufacturer.

### Circulating Oxidative Stress

The concentration of oxidized proteins in plasma samples was quantified using the OxiSelect Advanced Oxidation Protein Products (AOPP) Assay Kit (Cell Biolabs, Inc, Cat. No. STA-318). Plasma samples were diluted 1:2 in assay buffer prior to performing the assay according to manufacturer instructions with modifications. Briefly, diluted plasma samples were analyzed with and without (water added in equal volume) the reaction initiator in the same plate in order to calculate plasma background readings at an optical density of 340 (OD_340_). Plasma background values were then subtracted from the respective plasma OD_340_ before quantifying AOPP concentrations using reference to the standard curve.

### Circulating Corticosterone

Plasma corticosterone levels were assessed using a MILLIPLEX Hormone Magnetic Bead Panel (Millipore Sigma Cat. No. MSHMAG-21K). Plasma steroid hormones were extracted using acetonitrile extraction methods according to manufacturer instructions. Briefly, 375 µl of acetonitrile (Fisher Bioreagents, Cat. No. BP2405-1) was added to 250 µl of plasma sample to precipitate proteins. The precipitated solution was centrifuged at 17,000 x g for 10 minutes, and 500 µl of supernatant was collected and dried using vacuum centrifugation. Dried pellets were suspended in 200 µl of assay buffer and stored at −20⁰C. MILLIPLEX immunoassay was performed according to manufacturer instructions with an 18-hour overnight incubation. Samples, standards, and quality controls were plated in duplicate and corticosterone concentrations were measured on Luminex 200 instrument using xPONENT software version 4.3 (Luminex Corporation, Austin, TX). Quality control values for corticosterone were within the range provided by the manufacturer.

### Western Blot Analysis

Homogenized amygdala (basolateral and basomedial) and CA1 samples were denatured in Laemmli buffer (Bio-RAD, Cat. No. 161-0747) containing 10% β-mercaptoethanol (Fisher, Cat. No. BP176) and heated to 95°C for 5 minutes. Equal volumes of denatured samples containing 20 µg of total protein were loaded onto 4-15% polyacrylamide gels [Bio-RAD, Cat. No. 4561084 (CA1) or 4561086 (amygdala)] and resolved in tris-glycine running buffer (Bio-RAD, Cat. No. 1610771) at 25 mA for 1.5 hours at room temperature. Resolved proteins were transferred to PVDF membranes at 80 V for 2 h at 4°C. Membranes were blocked for 1 hour at room temperature with 5% non-fat milk in 1X tris-based saline-Tween-20 (TBST, Thermo Scientific, Cat. No. 28360). Membranes were incubated overnight at 4°C with primary antibodies probing Spectrin and enzyme-mediated degradation products (mouse anti-Spectrin, 1:1000, Millipore Sigma, Cat. No. MAB1622) and early growth response protein-1 (mouse anti-Egr-1, 1:100, Santa Cruz Biotechnology, Cat. No. sc-515830) diluted in 1% non-fat milk/TBST solutions. For protein normalization, we used β-actin primary antibody diluted in 1% non-fat milk/TBST solution (1: 3000, GeneTex, Cat. No. GTX629630) and incubated for 1 hour at room temperature. Membranes were washed in 10 minute increments in TBST for 30 minutes prior to incubation with secondary antibody. Membranes were incubated for 1 hour at room temperature with horseradish peroxidase-conjugated horse anti-mouse IgG solution (1:2500, Cell Signaling, Cat. No. 7074P2) diluted in 1% non-fat milk/TBST. Immunoreactive bands were visualized using a West Pico (Thermo Scientific, Cat. No. 34580) or West Femto (Thermo Scientific, Cat. No. 34095) enhanced chemiluminescence detection assay in a Syngene G:Box imager using GeneSys Image Acquisition software (Syngene, version 1.5.2.0). Band densitometry was quantified using Image J software (National Institutes of Health, version 1.53t). Total Spectrin (270 kD) and enzymatic cleavage of Spectrin by Calpain (150 kD) and Caspase-3 (120 kD) were first normalized to β-actin (42 kD) values before normalizing cleavage products to total Spectrin values. Egr-1 (54-58 kD) values were normalized to β-actin values. Quantification of protein expression is presented as % NULLI, where samples are normalized to NULLI samples of the same cohort ran in the same western blot. Representative images of full western blot membranes can be found in Supplemental Fig. S1.

### Data and Statistical Analyses

Data and statistical analyses were conducted in PRISM software (GraphPad, version 9.2). Outliers were identified and removed using robust regression and outlier removal (ROUT) with a coefficient Q = 1%. Data distributions were tested for normality using Shapiro-Wilk test. Data determined to have non-Gaussian distributions were log transformed before applying parametric statistics. One-Way ANOVA followed by Tukey’s multiple comparisons test was used to compare group differences unless otherwise noted. The relationship between inflammatory cytokines in the circulation and hippocampal CA1 was determined using Spearman correlation analysis. Spatial learning over the training period was assessed using Two-Way repeated measures ANOVA with Sidak’s multiple comparisons test. Spatial learning indices and memory were examined using unpaired t tests. The significance level was set to α = 0.05, and *p* ≤ 0.05 was considered significant. Values are expressed as means ± SD unless otherwise indicated.

## RESULTS

### Pregnancy-associated elevations in systemic inflammation are resolved during the post-pregnancy period and do not correlate with hippocampal inflammation during pregnancy

We observed elevated plasma pro-inflammatory (TNF-α, IL-6, IL-1β, IL-17A) and anti-inflammatory (IL-10) cytokines in pregnant rats during late gestation (GD21) compared to nulliparous rats (*p* ≤ 0.012, Fig. 1). Additionally, TNF-α and IL-10 plasma levels were reduced at two months post-pregnancy in primiparous rats compared to late gestation pregnant rats (*p* ≤ 0.017, Fig. 1). No significant differences were observed between nulliparous and primiparous rats in plasma levels of pro-inflammatory or anti-inflammatory cytokines (*p* > 0.05, Fig. 1). We next determined if pregnancy-associated elevations in circulating cytokines were reflected in dorsal hippocampal CA1, which is a brain region that contributes to cognitive function (43). There were no group differences in pro-inflammatory or anti-inflammatory cytokines in the CA1 (*p* > 0.05, Fig. 2). Moreover, we did not observe any correlations in circulating plasma cytokines with CA1 cytokine levels (*p* > 0.05, Supplemental Table S1).

**Fig. 1.**
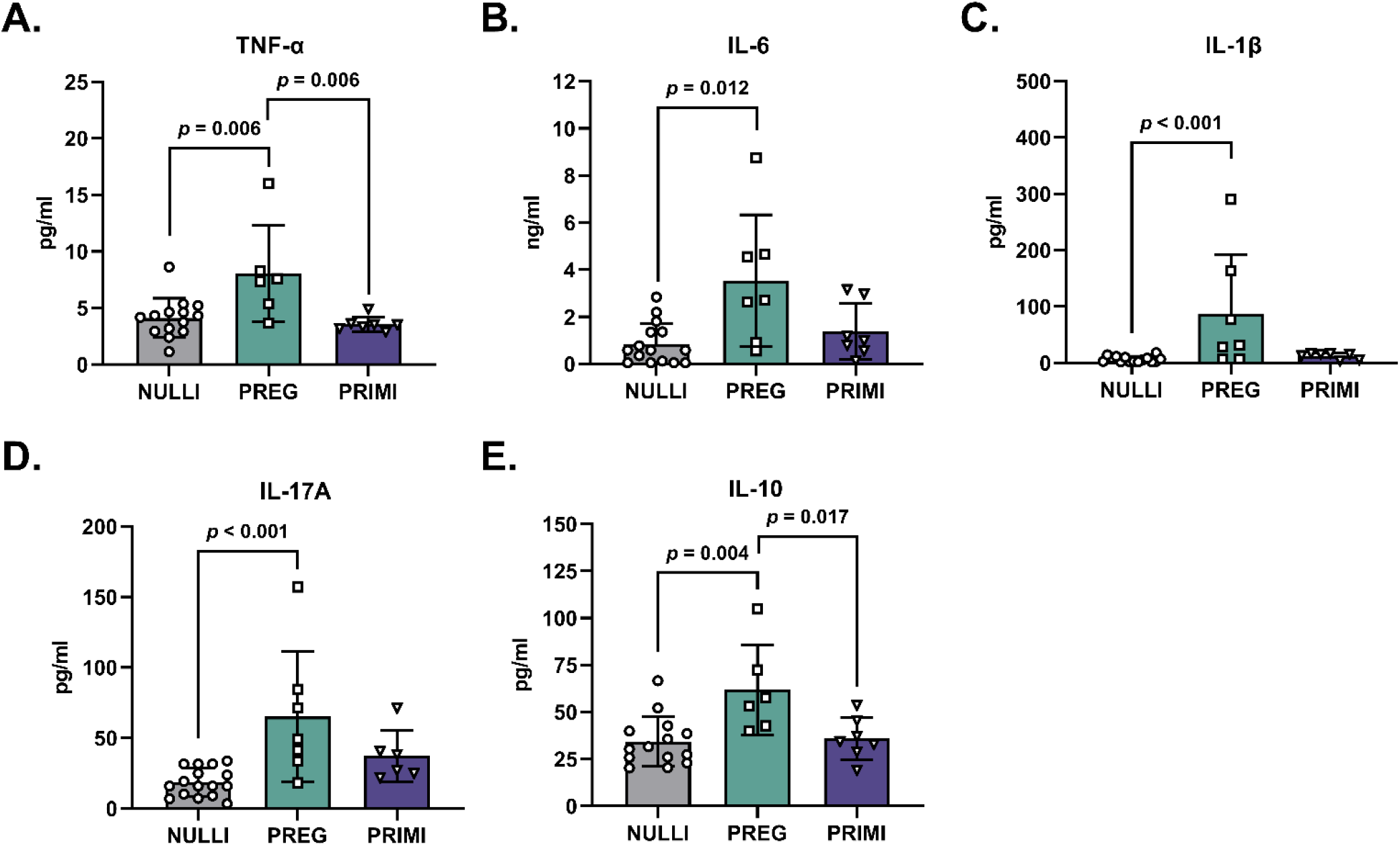
Circulating inflammatory cytokine profiles in healthy rats with various reproductive experiences. Plasma concentrations of pro-inflammatory (TNF-α, IL-6, IL-1β, IL-17A) and anti-inflammatory (IL-10) cytokines. NULLI = nulliparous (n = 14-16), PREG = gestational day 21 (n = 6-7), PRIMI = primiparous (two months post-pregnancy, n = 6-7). One-Way ANOVA with Tukey’s multiple comparisons test, mean ± SD.

**Fig. 2.**
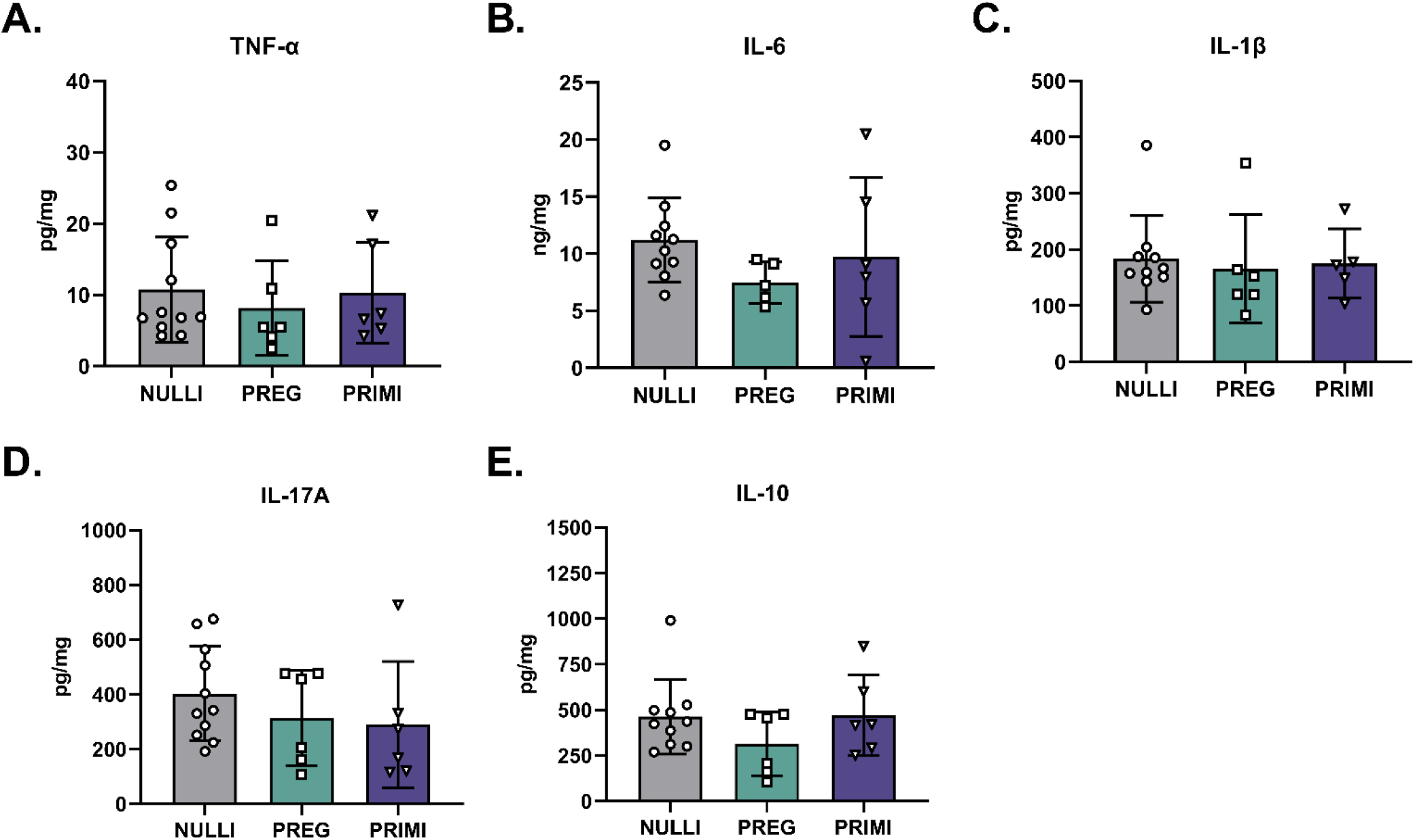
Inflammatory cytokine profiles in dorsal hippocampal CA1 of healthy rats with various reproductive experiences. Concentrations of pro-inflammatory (TNF-α, IL-6, IL-1β, IL-17A) and anti-inflammatory (IL-10) cytokines within the CA1 of the dorsal hippocampus. NULLI = nulliparous (n = 10-11), PREG = gestational day 21 (n = 5-6), PRIMI = primiparous (two months post-pregnancy, n = 5-6). One-Way ANOVA with Tukey’s multiple comparisons test, mean ± SD.

### Pregnancy is associated with elevated circulating and hippocampal oxidative stress

Advanced oxidation protein products were elevated in the plasma of pregnant rats compared to nulliparous and primiparous rats (*p* < 0.001, Fig. 3A). Additionally, oxidative stress-associated Calpain activation in the dorsal hippocampal CA1 was elevated in pregnant rats compared to nulliparous rats (*p* = 0.008, Fig. 3B and C). We then determined if there were any effects of pregnancy on markers of apoptosis and neuronal activation within the CA1. We did not observe any group differences in Caspase-3 activation (*p* > 0.05, Fig. 3B and D) or neuronal activation (*p* > 0.05, Fig. 3E and F) within the dorsal hippocampal CA1.

**Fig. 3.**
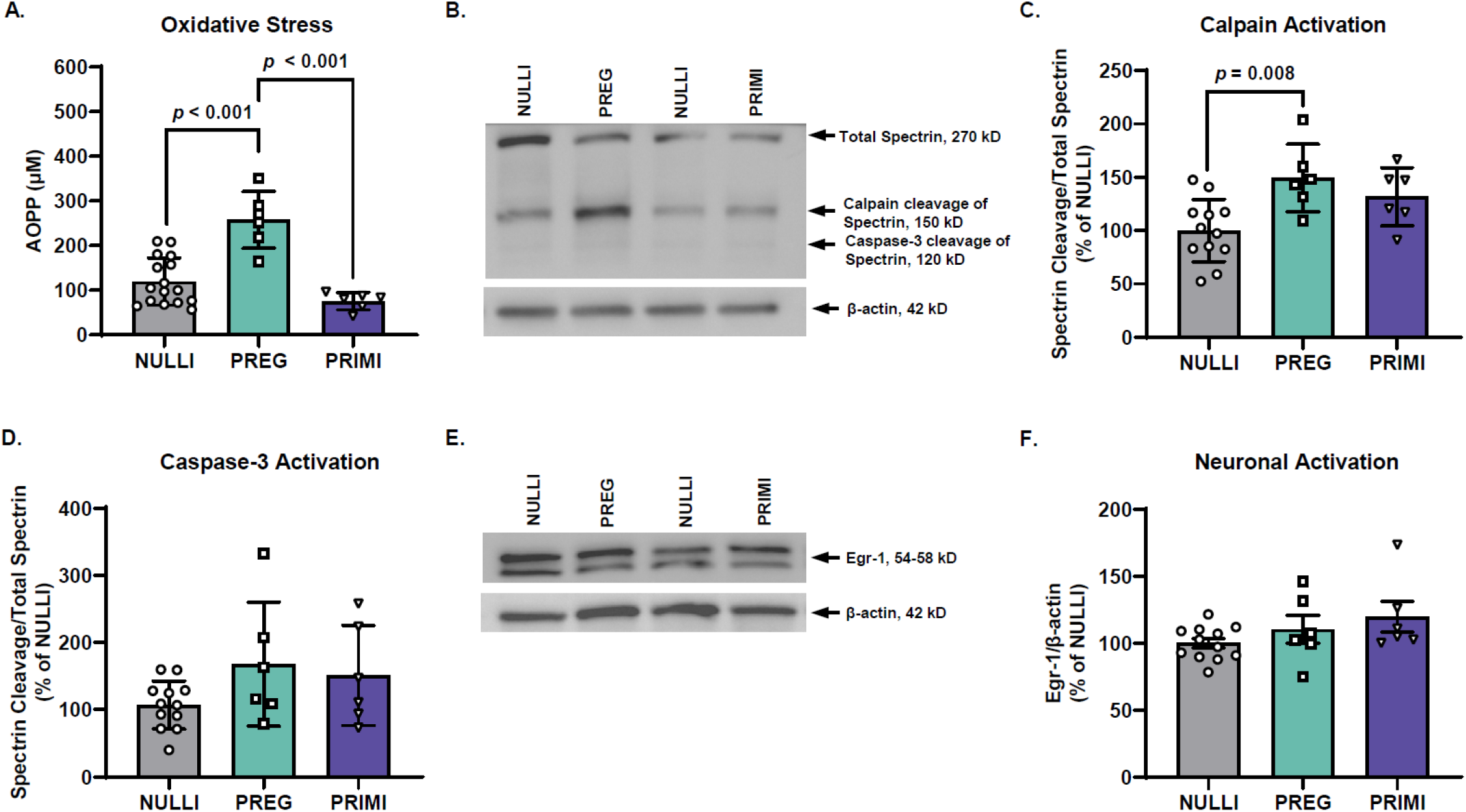
Circulating oxidative stress and cell activity in CA1 of dorsal hippocampus. **A)** Plasma concentrations of advanced oxidation protein products (AOPP, measure of oxidative stress). **B)** Western blot image of enzyme-mediated cleavage of Spectrin (270 kD), resulting in cleaved fragments at 150 kD (Calpain-mediated) and 120 kD (Caspase-3-mediated). **C)** Quantification of Calpain-mediated cleavage of Spectrin (measure of oxidative stress) in CA1. **D)** Quantification of Caspase 3-mediated cleavage of Spectrin (measure of apoptosis) in CA1. **E)** Protein expression of early growth response protein-1 (Egr-1, neuronal activation marker) in CA1. NULLI = nulliparous (n = 12-16), PREG = gestational day 21 (n = 6), PRIMI = primiparous (two months post-pregnancy, n = 6). Total Spectrin and cleavage products were first normalized to β-actin and a proportion of cleaved product to total Spectrin was analyzed. Protein expression shown as percentage of Nulliparous (NULLI). One-Way ANOVA with Tukey’s multiple comparisons test, mean ± SD.

### Primiparity does not impair recollective memory during pregnancy or post-pregnancy

Since we observed pregnancy-induced elevations in oxidative stress within the CA1 of pregnant rats, we determined if CA1-mediated cognitive performance was affected using the novel object recognition behavior task. Pregnant rats showed increased latency to initial contact with the novel object (*p* ≤ 0.004, Fig. 4A) and decreased total novel object contacts (*p* ≤ 0.045, Fig. 4B) compared to nulliparous and primiparous rats. However, pregnant rats also traveled less throughout the test compared to nulliparous and primiparous rats (*p* ≤ 0.005, Fig. 4C). Additionally, pregnant rats took longer to initiate movement in the behavior test compared to nulliparous rats (*p* = 0.004, Fig. 4D). Although there were no group differences in latency to initial contact of the familiar object (*p* > 0.05, Supplemental Fig. S2A), pregnant rats exhibited reduced total contacts with the familiar object compared to nulliparous and primiparous rats (*p* ≤ 0.025, Supplemental Fig. S2B). Furthermore, there were no group differences in object preference, as all groups spent more time investigating the novel object compared to time investigating the familiar object (*p* > 0.05, Supplemental Fig. S2C).

**Fig. 4.**
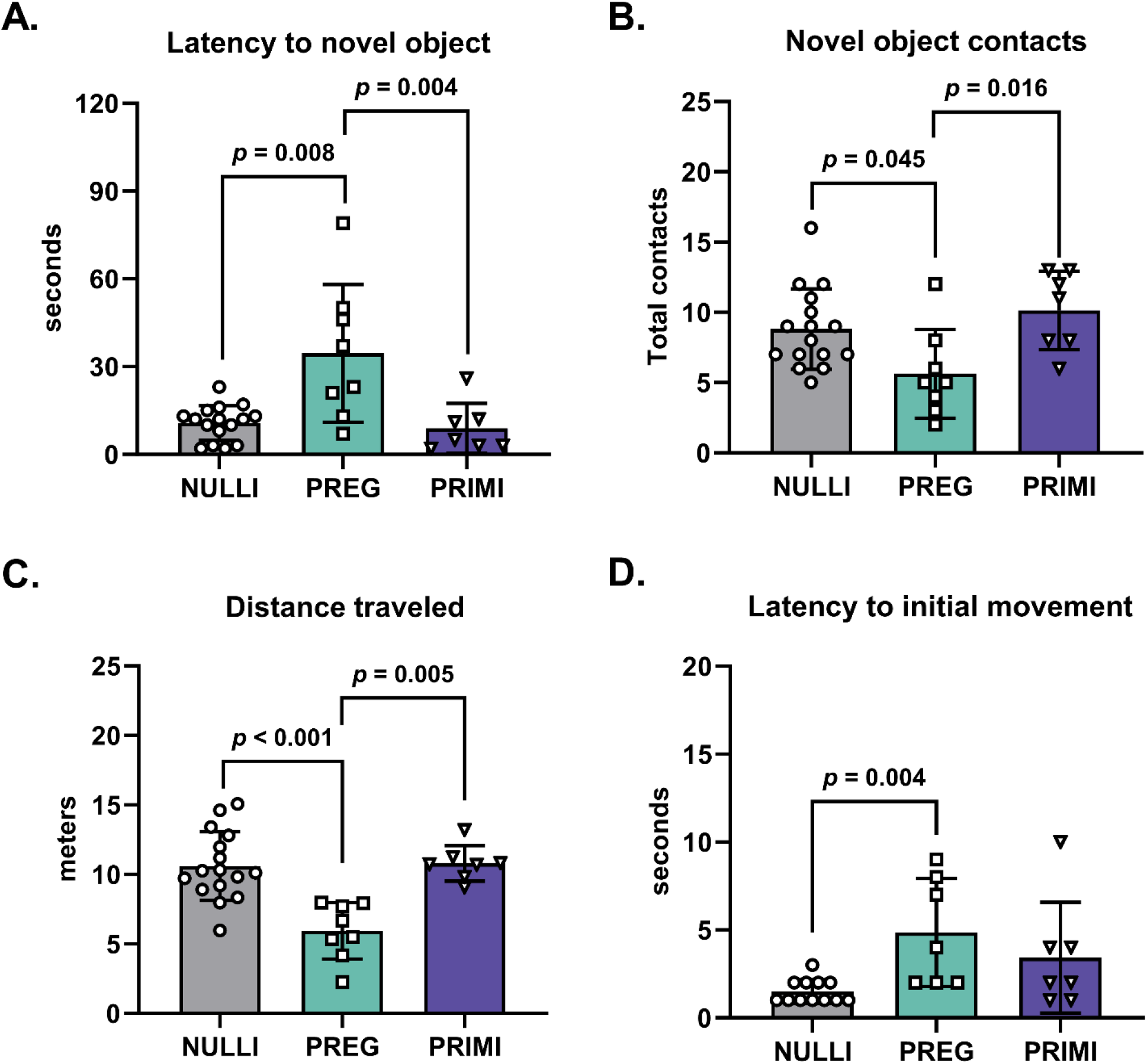
Recollective memory during novel object recognition behavior task. **A)** Latency to initial contact with novel object. **B)** Total contacts with novel object. **C)** Total distance traveled during test. **D)** Latency to initial movement during test. NULLI = nulliparous (n = 12-16), PREG = gestational day 20 (n = 7-8), PRIMI = primiparous (two months post-pregnancy, n = 7). One-Way ANOVA with Tukey’s multiple comparisons test, mean ± SD.

### Primiparity diminishes learning during the post-pregnancy period

To examine hippocampal-dependent spatial learning memory function, we performed the Morris water maze (Fig. 5) in age-matched nulliparous and primiparous rats one month before performing the novel object recognition test (Fig. 4). Due to additional stress of a swimming test, we did not test pregnant rats during late gestation in Morris water maze to prevent additional stressors during pregnancy that could alter pregnancy outcomes. We observed main effects of parity (*p* = 0.009) and learning day (*p* = 0.012) with no interactive effects (*p* = 0.768) on latency to target during learning days 2-4 (Fig. 5A). Additionally, primiparous rats exhibited increased latency to target compared to nulliparous rats on learning day 4 (*p* = 0.048, Fig. 5A). When assessing latency to target, nulliparous rats were better learners than primiparous rats during days 2-4 of learning (*p* = 0.007, Fig. 5B). However, no group differences were observed when assessing spatial memory on day 5 of the Morris water maze (*p* = 0.914, Fig. 5C). Similar to latency to target, we observed main effects of parity (*p* = 0.009) and learning day (*p* = 0.014) with no interactive effects (*p* = 0.371) on pathlength to target on learning days 2-4 (Fig. 5D). Nulliparous rats demonstrated shorter pathlengths to target on day 4 compared to day 2 of learning (*p* = 0.040), while primiparous rats did not exhibit shorter pathlengths as learning days progressed (*p* = 0.342, Fig. 5D). Similar to latency to target, nulliparous rats were better learners than primiparous rats when assessing pathlength to target (*p* = 0.001, Fig. 5E); yet, no group differences in spatial memory were observed on day 5 when evaluating pathlength to target (*p* = 0.372, Fig. 5F).

**Fig 5.**
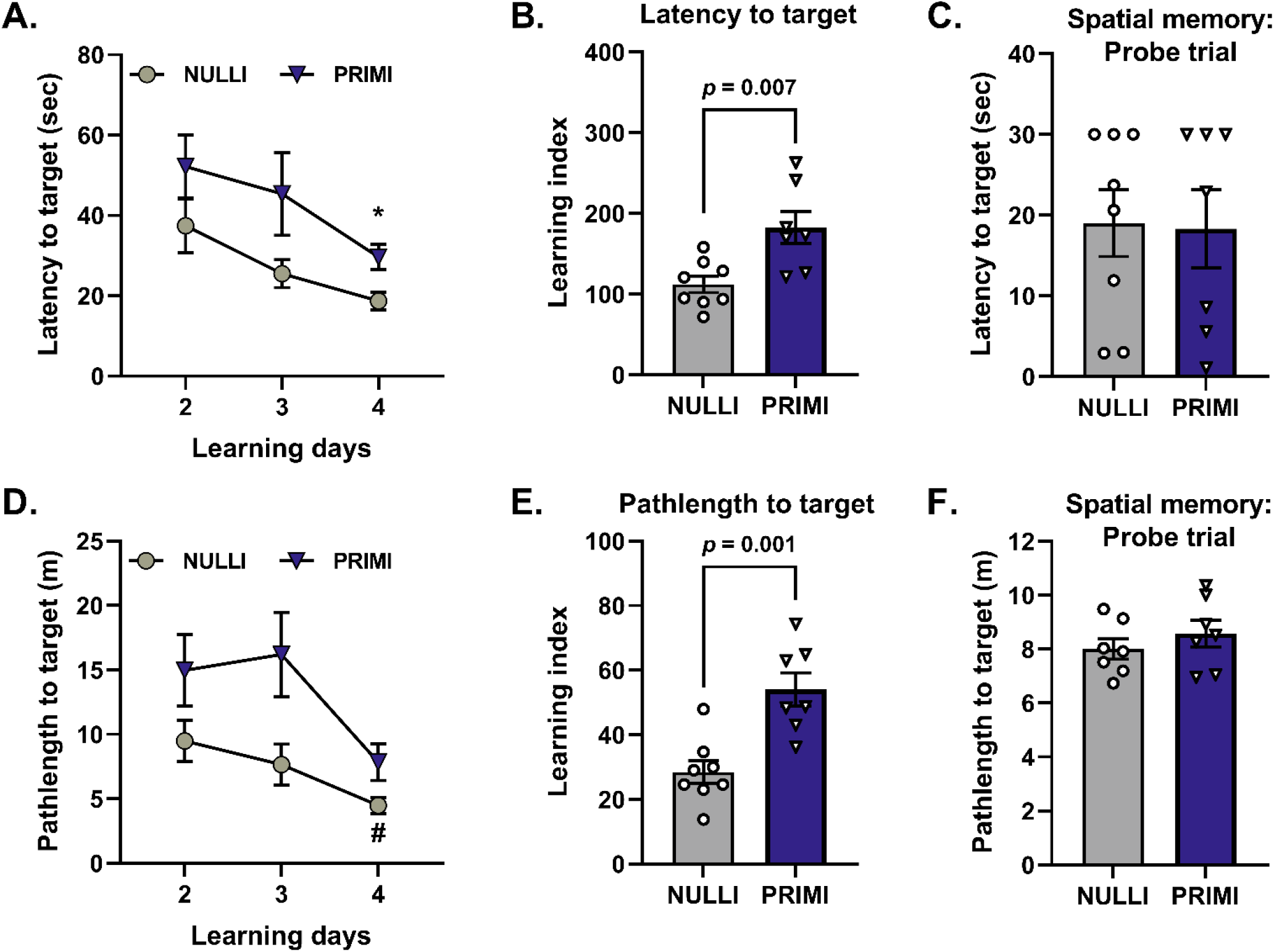
Learning indices and spatial memory during Morris water maze. **A)** Latency to target on learning days 2-4 of behavior task. **B)** Learning index calculated for each rat based on latency to target on learning days 2-4. A lower learning index represents better learning. **C)** Spatial memory assessed via latency to target on day 5 of behavior task. **D)** Pathlength to target on learning days 2-4 of behavior task. **E)** Learning index calculated for each rat based on pathlength to target on learning days 2-4. A lower learning index represents better learning. **F)** Spatial memory assessed via pathlength to target on day 5 of behavior task. NULLI = nulliparous (n = 7-8), PRIMI = primiparous [one week post-weaning (post-pregnancy day 35), n = 7]. Two-Way repeated measures ANOVA with Sidak’s multiple comparisons test **(A, D)** or unpaired t-test **(B, C, E, F)**, mean ± SEM. **\***, *p* < 0.05 vs PRIMI Day 4; **#**, *p* < 0.05 vs NULLI Day 2.

### Primiparity reduces anxiety-like behavior during the post-pregnancy period

Primiparous rats exhibited increased number of entries into the center of the arena compared to nulliparous and pregnant rats (*p* ≤ 0.034, Fig. 6A). However, primiparous rats were not spending more time in the center (*p* > 0.05, Fig. 6B) or traveling a greater distance in the center (*p* > 0.05, Fig. 6C) compared to nulliparous and pregnant rats. Yet, pregnant rats exhibited decreased distance traveled in the center compared to nulliparous rats (*p* = 0.012, Fig. 6C). Similar to the novel object recognition behavior task, pregnant rats traveled significantly less during the open field behavior test compared to nulliparous and primiparous rats (NULLI, mean ± SD: 16.53 ± 2.63 m, PREG: 12.62 ± 1.14 m, PRIMI: 19.60 ± 2.95 m, *p* ≤ 0.002). We observed no group differences in circulating levels of the stress hormone corticosterone (*p* > 0.05, Fig. 6D). Additionally, we observed no differences in the oxidative stress marker, Calpain-mediated Spectrin cleavage, in the amygdala [p > 0.05; (% NULLI, mean ± SD), NULLI: 100.0 ± 32.19%, PREG: 108.9 ± 42.73 %, PRIMI: 133.0 ± 69.01 %, Supplemental Fig. S1].

**Fig. 6.**
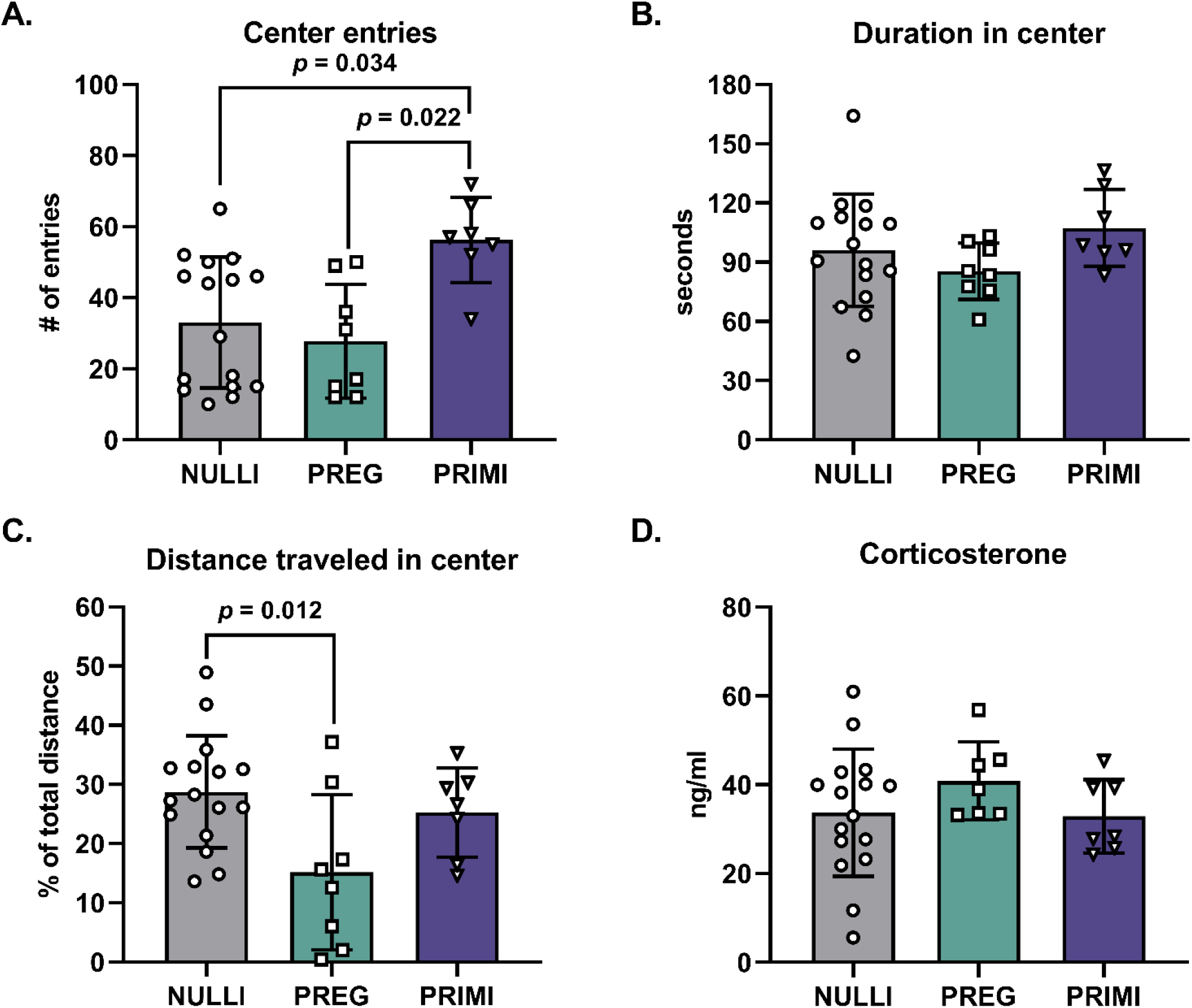
Anxiety-associated behavior during open field behavior task. **A)** Total entries into the center of the open arena. **B)** Duration in seconds in center of open area. **C)** Distance traveled in the center of the open arena divided by the total distance traveled during behavior task. **D)** Plasma concentrations of stress-associated steroid hormone corticosterone. NULLI = nulliparous (n = 16), PREG = gestational day 20 (n = 7-8), PRIMI = primiparous (two months post-pregnancy, n = 7). One-Way ANOVA with Tukey’s multiple comparisons, mean ± SD.

## DISCUSSION

In this study, we determined the impact of reproductive status on systemic and localized inflammation and oxidative stress, as well as cognitive performance in rats. We found that systemic inflammation and oxidative stress are elevated during healthy rodent pregnancy but resolve in the post-pregnancy period. In addition, maternal oxidative stress-associated enzymatic activity was elevated in the maternal dorsal hippocampal CA1 during pregnancy while inflammatory cytokines remained unchanged. Contrary to systemic oxidative stress, maternal CA1 oxidative stress levels were comparable in pregnant and primiparous rats at two month post-pregnancy. When assessing cognitive performance, we found that reproductive status did not impact recollective or spatial memory during pregnancy or post-pregnancy. Although we did not observe memory impairments, we found that primiparous rats exhibit learning deficits and enhanced exploratory behavior in the center of an open arena in the post-pregnancy period.

This is the first study to examine the effects of reproductive experience on the association between maternal systemic and localized brain inflammatory cytokines. We and others have observed increases in systemic inflammation during late gestation of rodent pregnancy (44, 45). Few studies have examined cytokine levels during post-pregnancy and within the maternal brain during pregnancy and post-pregnancy (46–49). Here, we found that inflammatory cytokine levels within the dorsal hippocampal CA1 were similar among rats of varying reproductive experience while plasma inflammatory cytokines drastically increased during pregnancy, suggesting that the maternal CA1 is protected from pregnancy-associated elevations in systemic inflammation. These findings are consistent with previous reports demonstrating no changes in maternal hippocampal IL-6 or IL-1β during late gestation or early post-pregnancy in Sprague-Dawley rats (48). Similarly, a recent study by Duarte-Guterman et al. observed no effect of reproductive experience (nulliparity, primiparity, biparity) on hippocampal inflammatory cytokine production at 30 or 240 days post-pregnancy (47). Contrarily, a previous study by Haim et al. revealed increases in IL-6 and IL-10 within the maternal dorsal hippocampus at eight days post-pregnancy in primiparous rats (46). Notably, these previous studies examined cytokine expression within the whole dorsal hippocampus while the present study quantified cytokine expression specifically within the CA1 region of the dorsal hippocampus. Moreover, the current study and previous studies examined cytokine levels within the hippocampus at different reproductive states (i.e. late gestation or post-pregnancy days 8, 30, 60, and 240). Taken together, reproductive experience may differentially impact specific regions of the maternal dorsal hippocampus (e.g. CA1, dentate gyrus, CA3) depending on reproductive status (early pregnancy vs late pregnancy and early post-pregnancy vs late post-pregnancy). Future studies examining cytokine expression levels in other hippocampal-associated brain areas, such as the amygdala and entorhinal cortex, during pregnancy and post-pregnancy are warranted. Importantly, our findings of no correlations between systemic and localized cytokine expression levels highlights the restriction of pregnancy-associated systemic inflammation from infiltrating the maternal brain during pregnancy, as well as the limitation of plasma sampling as a proxy for maternal CA1 inflammation status.

This is also the first study to examine maternal systemic and hippocampal CA1 markers of oxidative stress during pregnancy and post-pregnancy. We found that oxidative stress markers are elevated within the plasma (AOPP) and maternal CA1 (Calpain enzymatic activity) during pregnancy. Importantly, systemic levels of oxidative stress declined in the post-pregnancy period while CA1 oxidative stress levels were comparable to levels observed during pregnancy, suggesting that pregnancy-associated elevations in oxidative stress may be longer-lasting in maternal tissues in comparison to the maternal circulation. Although oxidative stress-associated enzymatic activity was elevated within the maternal CA1 during pregnancy, we did not observe evidence of cell death (Caspase-3 activation) or decreased neuronal activation. Collectively, the observed elevations in oxidative stress-associated markers occurred as a physiological response during healthy rodent pregnancy and suggest that the maternal CA1 is vulnerable to oxidative stressors during pregnancy; nonetheless, physiological elevations in maternal CA1 oxidative stress does not elicit neuronal damage within this brain region or hippocampal-associated cognitive impairment. Similar to studies in maternal brain inflammation, future studies are needed to assess the impact of maternal oxidative stress in various brain regions associated with maternal behavior. Currently, there is growing evidence supporting the impact of pathophysiological elevations in oxidative stress during pregnancy, such as in gestational disorders, on fetal and offspring measures of brain oxidative stress and cognitive function (50). Yet, there is a paucity of studies examining the effect of elevated pregnancy-associated oxidative stress on maternal brain oxidative stress and cognitive function. Clinically, women who experience pregnancy complications associated with elevated oxidative stress, such as hypertensive disorders of pregnancy, are more likely to self-report cognitive impairments and have cognitive decline (19–21). Future studies interrogating oxidative stress within the maternal brain are needed to understand the role of maternal brain oxidative stress in physiological and pathophysiological reproductive states, including in animal models of gestational diabetes, hypertensive disorders of pregnancy, and maternal obesity.

When assessing the impact of reproductive experience on recollective memory in the novel object recognition behavior task, pregnant rats exhibited increased latency to initial contact with the novel object and reduced overall novel object contacts. However, there were no differences among the groups in the preference for the novel object. Of note, pregnant rats traveled less during the behavior task and were slower to initiate movement when compared to both nulliparous and primiparous rats, which likely contributed to latency to the first interaction with the novel object and the total number of contacts over the test duration. Altogether, our data suggest that reproductive experience does not impact recollective memory. This finding is in agreement with previous observations examining recollective memory in nulliparous, primiparous, and multiparous Long Evans rats (51). Our findings also highlight the limitation of using the novel object recognition behavior task for assessing recollective memory in pregnant rats. Given pregnant rats exhibit reduced locomotion, especially during late gestation, future studies could incorporate memory tasks that reduce mobility constraints, such as home cage-based recognition tasks that limit the arena size (52).

Similar to our findings on recollective memory, we also observed no effect of reproductive experience on spatial memory when assessing latency and pathlength to target during the probe trial of the Morris water maze. However, we did observe learning impairments in primiparous rats compared to nulliparous rats, as evidenced by increases in the learning indices for latency to target and pathlength to target over the learning period (days 2-4). These findings in primiparous rats 35 days post-pregnancy extend previous studies that demonstrated impaired spatial learning in the Morris water maze in primiparous rats 1-5 days post-pregnancy (53). Of note, we incorporated the Morris water maze one month prior to the novel object recognition behavior task in our experimental design to prevent confounds associated with battery testing (54, 55). Additionally, we conducted the Morris water maze one week following weaning to also prevent any confounds related to weaning and lactation. Based on prior reports indicating the potential of the Morris water maze to induce stress during rodent pregnancy (56), we did not examine spatial memory in pregnant rats. Nevertheless, previous studies have demonstrated that pregnant rats exhibit poorer spatial learning in the Morris water maze compared to nulliparous rats (56). It is postulated that impaired spatial learning is an adaptive response associated with nesting and preparing for parturition (56), and may extend into the post-pregnancy period as mothers care for their young. Indeed, our findings reveal that spatial learning remains impaired at 35 days post-pregnancy. Interestingly, a previous study by Pawluski et al. revealed spatial learning and memory are enhanced at 55 days post-pregnancy compared to nulliparous rats (57), suggesting that previous pregnancy experience and time since pup separation have an impact on maternal spatial learning and memory.

Since pregnant rats were delayed in initial movement and demonstrated decreased total distance traveled during the novel object recognition behavior task, we examined anxiety-like behavior in an open field arena. Our findings demonstrate that primiparous rats exhibit reduced anxiety-like behavior in the open field behavior task, which is consistent with previous findings in young primiparous rats (58). However, primiparous rats did not spend more time or travel a greater distance in the center in the present study, revealing that center entries were abrupt. Contrarily, pregnant rats exhibited fewer center entries and shorter distances traveled in the center. Although this may be perceived as an anxiety-like behavior, we caution this interpretation given pregnant rats also traveled less throughout the behavior task compared to nulliparous and primiparous rats, as we observed similarly in the novel object behavior task.

Due to group differences observed in anxiety-like behaviors, we investigated circulating stress hormones and oxidative stress within the amygdala, a brain region associated with anxiety-like behavior (59). We did not observe an impact of reproductive experience on plasma levels of the stress hormone corticosterone, which agrees with previous studies that examined corticosterone levels in primiparous rats during pregnancy and post-pregnancy (60, 61). To our knowledge, this is the first study examining oxidative stress and cell death-associated enzymatic activity within the maternal amygdala. Our findings indicate that there were no group differences in oxidative stress-associated enzymatic activity or cell death within the amygdala. This is in contrast with our oxidative stress-associated observations in the CA1 during pregnancy, which further highlights the effects of reproductive experience on maternal brain oxidative stress to be brain region-specific.

### Limitations, Perspectives, and Significance

Our study has numerous strengths including assaying systemic and localized cytokines and oxidative stress measures, conducting behavioral analyses during the rodent active cycle and outside of weaning in primiparous rats to prevent pup separation and lactation-associated hormonal confounds, including complementary behavior tests to determine if there were any pregnancy-associated confounds, and assessing hippocampal-associated maternal behaviors alongside region-specific molecular analyses. Even so, there are several limitations in this study that warrant further analyses. For instance, we did not assess the impact of reproductive experience on blood-brain barrier permeability, which may contribute to vulnerability of brain-specific regions to oxidative stressors during physiological and pathophysiological pregnancies (62). Additionally, the reduced locomotive activity in pregnant rats during late gestation warrants the modification of memory and anxiety-like behavior tasks to accommodate reductions in overall movement. Lastly, we did not examine the impact of reproductive experience on specific cell types within the maternal CA1 that could be affected by systemic and local elevations in oxidative stress and inflammation. Previous studies have revealed reductions in the density of brain-resident macrophages, microglia, during pregnancy and early postpartum (46, 49), but interactions of microglia with neurons and their role in modulating maternal behavior remains unclear. Future studies could be aimed at understanding the impact of reproductive experience and oxidative stress on cell-cell interactions within various maternal brain regions during pregnancy and post-pregnancy. Finally, our experimental design focused on late term and two months post-pregnancy, in which we observed elevations in oxidative stress within the maternal CA1. Future studies could incorporate temporal oxidative stress analyses spanning pregnancy and post-pregnancy to provide a better understanding of fluctuations in oxidative stress within the maternal brain.

In conclusion, oxidative stress-associated markers, and not inflammatory cytokines, were elevated in the maternal CA1 during healthy pregnancy, revealing a vulnerability of the maternal hippocampal CA1 to oxidative stressors. Even so, elevations in physiological oxidative stress within the maternal brain did not impact neuronal activity or cognitive performance during pregnancy. However, a previous healthy pregnancy history resulted in rodent learning deficits and reduced anxiety-like behavior, highlighting long-term effects of healthy pregnancy on maternal cognitive function. Future studies examining pathophysiological elevations in maternal brain oxidative stress, such as in pregnancy complications with hypoxic insults, may reveal underlying mechanisms contributing to associated adverse neural and behavioral plasticity in the transition to motherhood.

## DATA AVAILABILITY

Data will be made available upon reasonable request.

## GRANTS

This study was supported by National Institutes of Health Grant R01 HL146562-04S1 (S.G.) and American Heart Association Grants 22POST-903250 (to J.L.B.), 22PRE-900431 (to J.J.G.), and 23PRE-1012811 (to S.M.T). The content is solely the responsibility of the authors and does not necessarily represent the official views of the National Institutes of Health.

## AUTHOR CONTRIBUTIONS

J.L.B., S.G., and R.L.C. conceived and designed research; J.L.B., E.N.W., J.J.G., S.M., S.M.T., and N.R. performed experiments; J.L.B., E.N.W, and E.V. analyzed data; J.L.B., S.G., and R.L.C. interpreted results of experiments; J.L.B. prepared figures; J.L.B. drafted manuscript; J.L.B., E.N.W., J.J.G., S.M., S.M.T., N.R., E.V., S.G., and R.L.C. edited and revised manuscript; J.L.B., E.N.W., J.J.G., S.M., S.M.T., N.R., E.V., S.G., and R.L.C. approved final version of manuscript.

