## Supplemental Material for "Pregnancy-associated oxidative stress and inflammation are not associated with impaired maternal neuronal activity or memory function"

**Table S1. Correlations of inflammatory cytokines in plasma and CA1 of dorsal hippocampus.**

| | **All groups (n = 21-23)** | | **NULLI (n = 9-12)** | | **PREG (n = 5-6)** | | **PRIMI (n = 5-6)** | |
|---|---|---|---|---|---|---|---|---|
| **Cytokines** | **r** | ***p*-value** | **r** | ***p*-value** | **r** | ***p*-value** | **r** | ***p*-value** |
| **TNF-α** | 0.41 | 0.52 | 0.51 | 0.09 | -0.10 | 0.95 | -0.06 | 0.93 |
| **IL-6** | 0.06 | 0.79 | 0.26 | 0.41 | -0.40 | 0.52 | 0.49 | 0.36 |
| **IL-1β** | -0.19 | 0.39 | 0.32 | 0.40 | -0.56 | 0.40 | -0.50 | 0.45 |
| **IL-17A** | -0.04 | 0.86 | 0.07 | 0.83 | 0.30 | 0.68 | 0.30 | 0.68 |
| **IL-10** | -0.07 | 0.75 | 0.23 | 0.47 | 0.10 | 0.95 | -0.77 | 0.10 |

**r** = Spearman's rank correlation coefficient. NULLI = nulliparous, PREG = gestational day 21, PRIMI = primiparous (two months post-pregnancy).

### Supplemental Figures

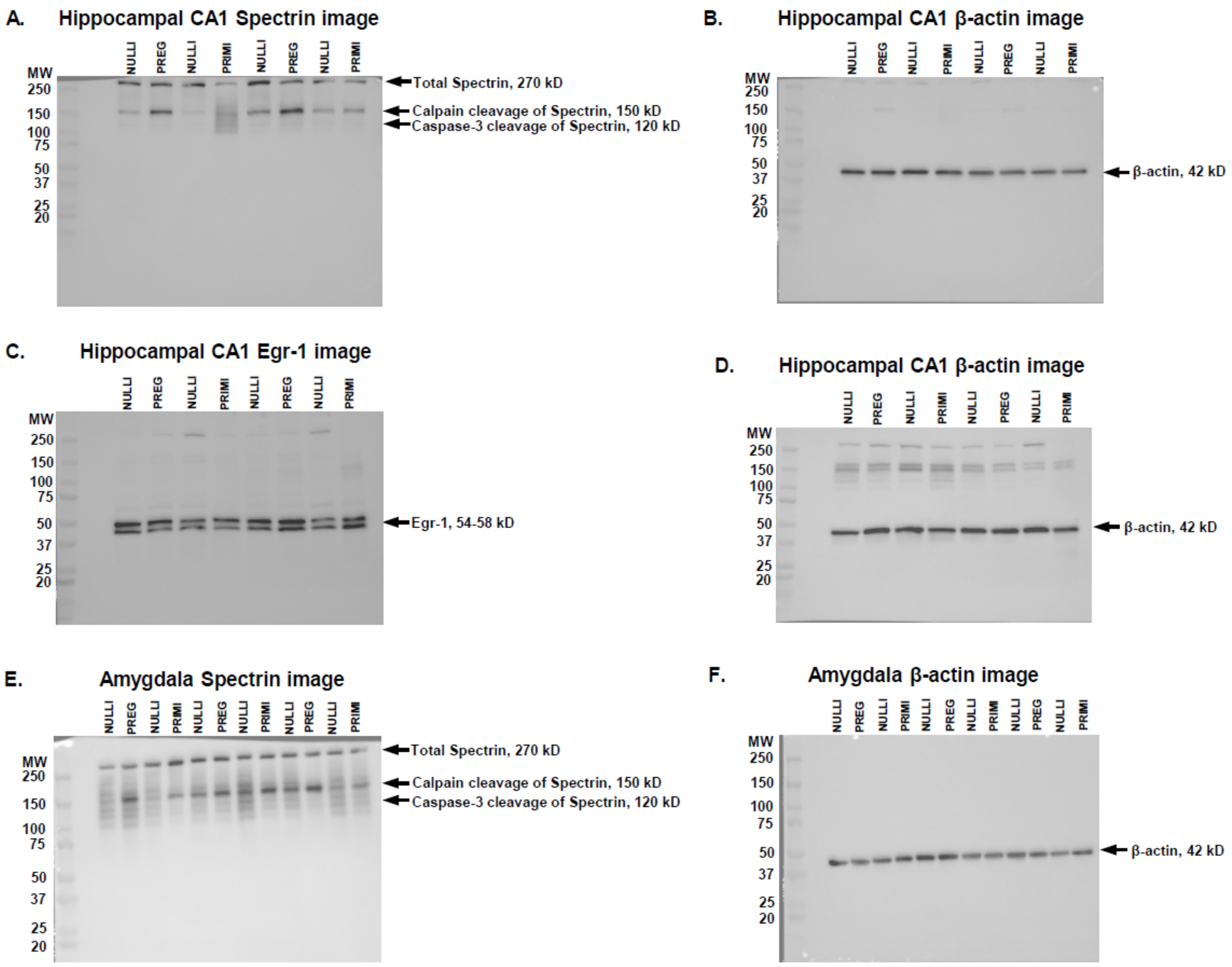


**Figure S1. Representative western blot images. A)** Spectrin protein expression and enzymatic cleavage by Calpain and Caspase-3 in dorsal hippocampal CA1. **B)** β-actin expression used for normalization of protein expression in **A**. **C)** Egr-1 protein expression in dorsal hippocampal CA1 normalized to β-actin expression **(D)**. **E)** Spectrin protein expression and enzymatic cleavage by Calpain and Caspase-3 in amygdala and normalized to β-actin **(F)**. Egr-1 = early growth response-1, MW = molecular weight, NULLI = nulliparous, PREG = gestational day 21, PRIMI = primiparous (two months post-pregnancy).

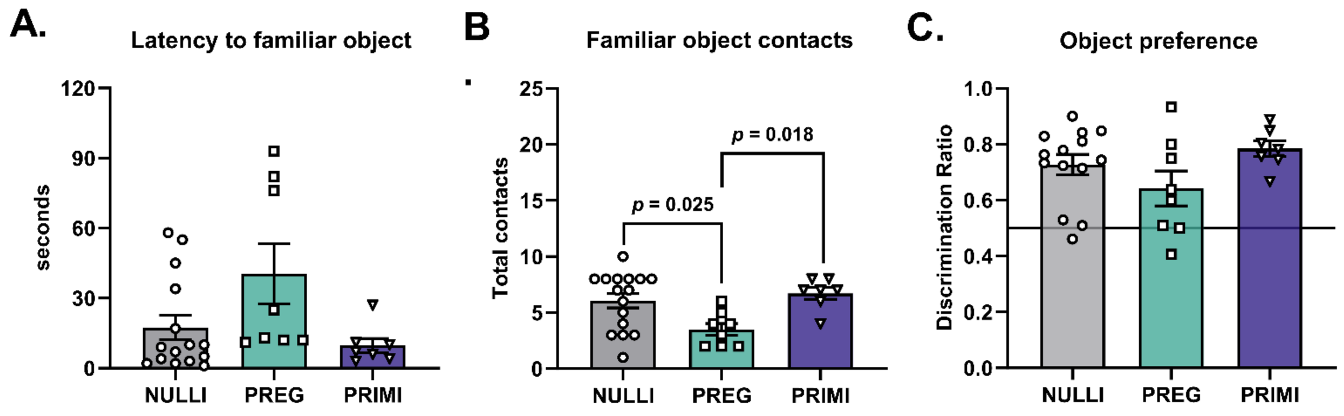


**Figure S2. Familiar object recognition and object preference during novel object behavior task. A)** Latency to initial contact with familiar object. **B)** Total contacts with familiar object. **C)** Discrimination ratio**,** defined as the duration in seconds spent exploring novel object divided by the duration exploring novel and familiar objects. Dark horizontal bar at y = 0.5 represents non-discrimination (equal time exploring novel and familiar object). NULLI = nulliparous (n = 14-16), PREG = gestational day 20 (n = 8), PRIMI = primiparous (two months post-pregnancy, n = 7). One-Way ANOVA with Tukey's multiple comparisons test, mean ± SD.
